# Neuronal Morphology Enhances Robustness to Perturbations of Channel Densities

**DOI:** 10.1101/2022.10.14.512316

**Authors:** Yunliang Zang, Eve Marder

## Abstract

Biological neurons show significant cell-to-cell variability but have the striking ability to maintain their key firing properties in the face of unpredictable perturbations and stochastic noise. Using a population of multi-compartment models consisting of soma, neurites, and axon for the lateral pyloric (LP) neuron in the crab stomatogastric ganglion, we explored how rebound bursting is preserved when the 14 channel conductances in each model are all randomly varied. The soma-axon coupling is critical for the ability of the axon to spike during bursts and consequently determines the set of successful solutions. When the coupling deviates from a biologically realistic range, the neuronal tolerance of conductance variations is significantly lessened. Thus, the gross morphological features of these neurons enhance their robustness to perturbations of channel densities and expands the space of individual variability that can maintain a desired output pattern.

## Introduction

It is now well-established that multiple sets of maximal conductances of ion channels can produce very similar neuronal activity patterns in both individual neurons and circuits (1-9). Nonetheless, it is also clear that models of desired intrinsic properties are relatively rarely found by randomly sampling within a space of parameters (5, 6). Thus, it is likely that biological neurons use homeostatic and other developmental mechanisms to successfully find rare sets of values of ion channel conductances that give rise to specific patterns of intrinsic excitability (10-14).

Although single-compartment, isopotential, conductance-based models have been enormously useful in understanding the relationships among ion channels that give rise to electrical excitability, most biological neurons have complex structures, with differential distributions of channels in the membranes of different neuronal regions. While there are specific examples of conductance-based models of complex neurons (15-20), these models have often been finely tuned to achieve their desired activity patterns. Thus, it is less clear in these more complex models the extent to which complex structures aid or diminish the likelihood of finding vastly different degenerate sets of conductance values that support characteristic activity patterns.

In this study, we exploit a previously developed 4-compartment model database of the Lateral Pyloric (LP) neuron of the crustacean stomatogastric ganglion (STG) to look at the influence of LP neuron structure on its robustness to perturbations of its channel densities. This is partly motivated by recent studies that show that the detailed neuropilar projections of the LP neuron have relatively little impact on how slow synaptic events are processed because of the large length constants of the major neurites (21-23). In these 4-compartment models (soma, axon, and two neurite compartments) only the axon compartment has voltage-gated Na^+^ channels (5). To create this model database, Taylor et al (5) generated ∼ 600,000 randomly selected sets of conductance densities and then sorted them to find ∼1,300 models with the desired firing patterns matching the biological properties of the LP neuron. Specifically, when isolated, the biological LP neuron fires tonically, but shows robust rebound bursts in response to inhibitory inputs from its presynaptic partners. These rebound bursts are important in ensuring that the LP neuron fires at an appropriate time during the triphasic pyloric rhythm (24).

The successful ∼1,300 models vary considerably in their maximal conductances and are examples of degenerate or multiple solutions that produce a characteristic behavior. In this paper, we look at the effect of neuronal structure on the robustness of this population of LP neuron models in response to perturbations of conductance densities.

## Results

Figure 1A shows a diagram of the chemical synaptic inputs to the LP neuron. The top trace in Figure 1B is an intracellular recording from the LP neuron, with its sequential inhibition from the PY (Pyloric), PD (Pyloric Dilator) and AB (Anterior Burster) neurons (note that AB and PD slow waves and spikes are synchronized), followed by the rebound burst in the LP neuron. The timing of the inhibitory inputs is shown by the extracellular recordings from the lvn (lateral ventricular nerve, Figure 1B, bottom). The LP neuron somatic action potentials that ride on the top of the slow wave are highly attenuated by the LP neuron’s cable, as they are generated in the distant axon.

**Fig. 1.**
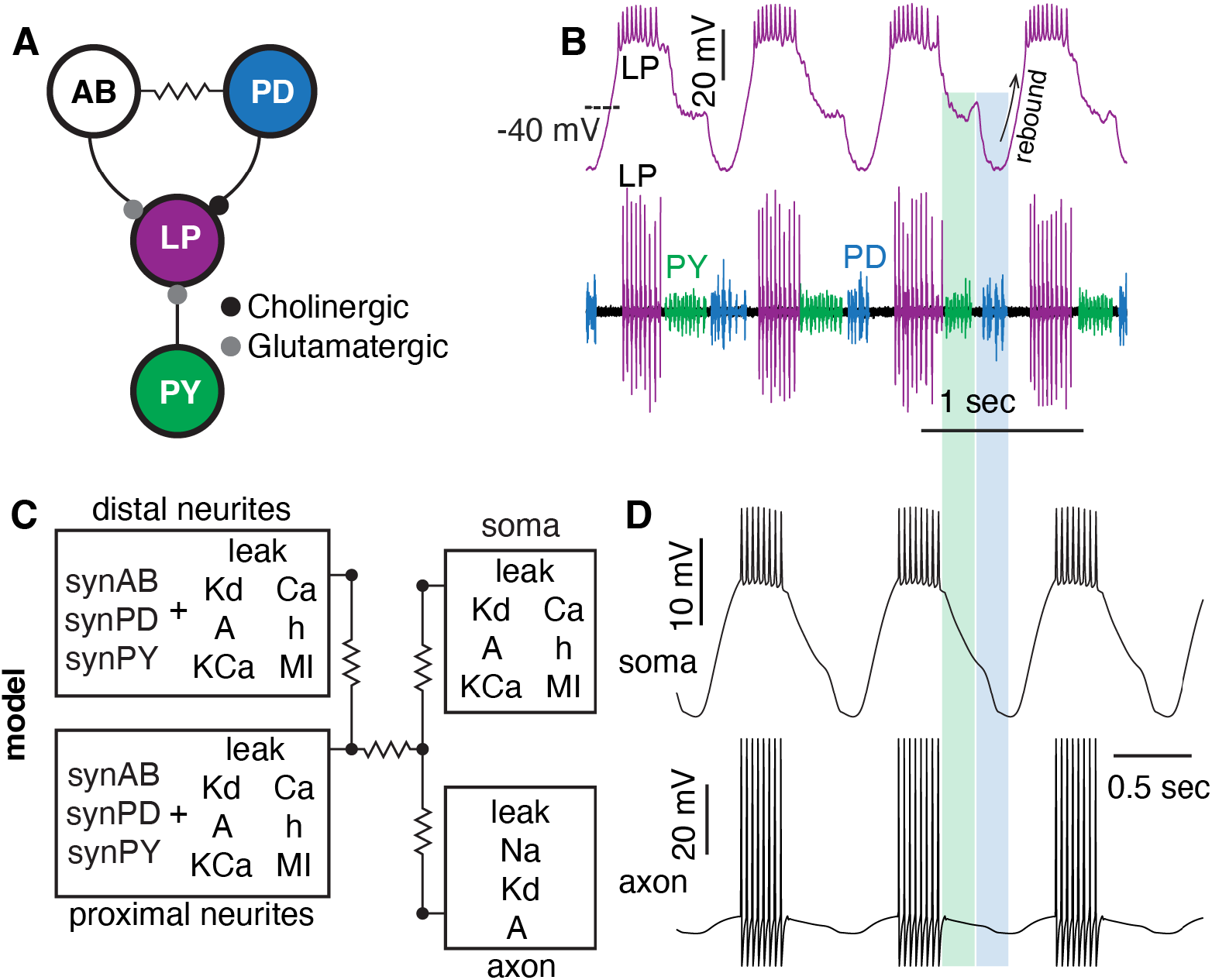
LP neuron multi-compartment model structure and spike properties. (*A*) Schematic of the chemical synaptic inhibition that a LP neuron receives. (*B*) Intracellular recordings of a LP neuron (top) and an extracellular recording at the lvn (bottom, spikes of LP, PY and PD neurons are color coded, unpublished data from Sonal Kedia). (*C*) Schematic of the LP neuron models, consisting of an axon, a soma, proximal neurites, and distal neurites with corresponding ion channels and synapses distributed in each compartment, modified from (5). (*D*) Example somatic (top) and axonal (bottom) spike waveforms in a model. The green and blue shadings illustrate the timing of the inhibition from PY neurons and PD neurons (also AB neuron).

Figure 1C shows the composition of ion channels in each of the 4 compartments in the model. The soma has a leak current (I_leak_), a modulator current (I_MI_), three potassium currents (I_Kd_, delayed-rectifier K^+^ current; I_A_, A-type K^+^ current; I_KCa_, calcium-activated K^+^ current), a calcium current (I_Ca_) and a hyperpolarization-activated inward current (I_h_). In addition to these currents, three inhibitory synaptic currents, from the AB, PD, and PY neurons (synAB, synPD and synPY) are also distributed in the neurites. The axon has a I_leak_, I_Kd_, I_A_, and a voltage-gated Na^+^ current (I_Na_). A characteristic waveform from all LP neuron models is seen in Figure 1D. The top trace is the model behavior in the soma compartment, while the bottom trace is the model behavior in the axon. The axon generates overshooting action potentials, and only a very small slow wave is visible, while the soma compartment replicates the somatic recordings seen in the biological neurons (Fig. 1B).

### Neuronal Robustness to Channel Conductance Variations

We used 1292 of Taylor’s LP neuron models (5), hereafter referred to as “parent” models whose parameters show large cell-to-cell variations despite similar output patterns. We first varied all of the 14 channel conductances at the same time by different random values in each “parent” model, based on their range of values for all of the 1292 models. Then, for each “parent” model, multiple runs, with different sets of channel perturbations, we generated a number of “child” models, which displayed a variety of intrinsic patterns of activity (Fig. 2A; methods).

**Fig. 2.**
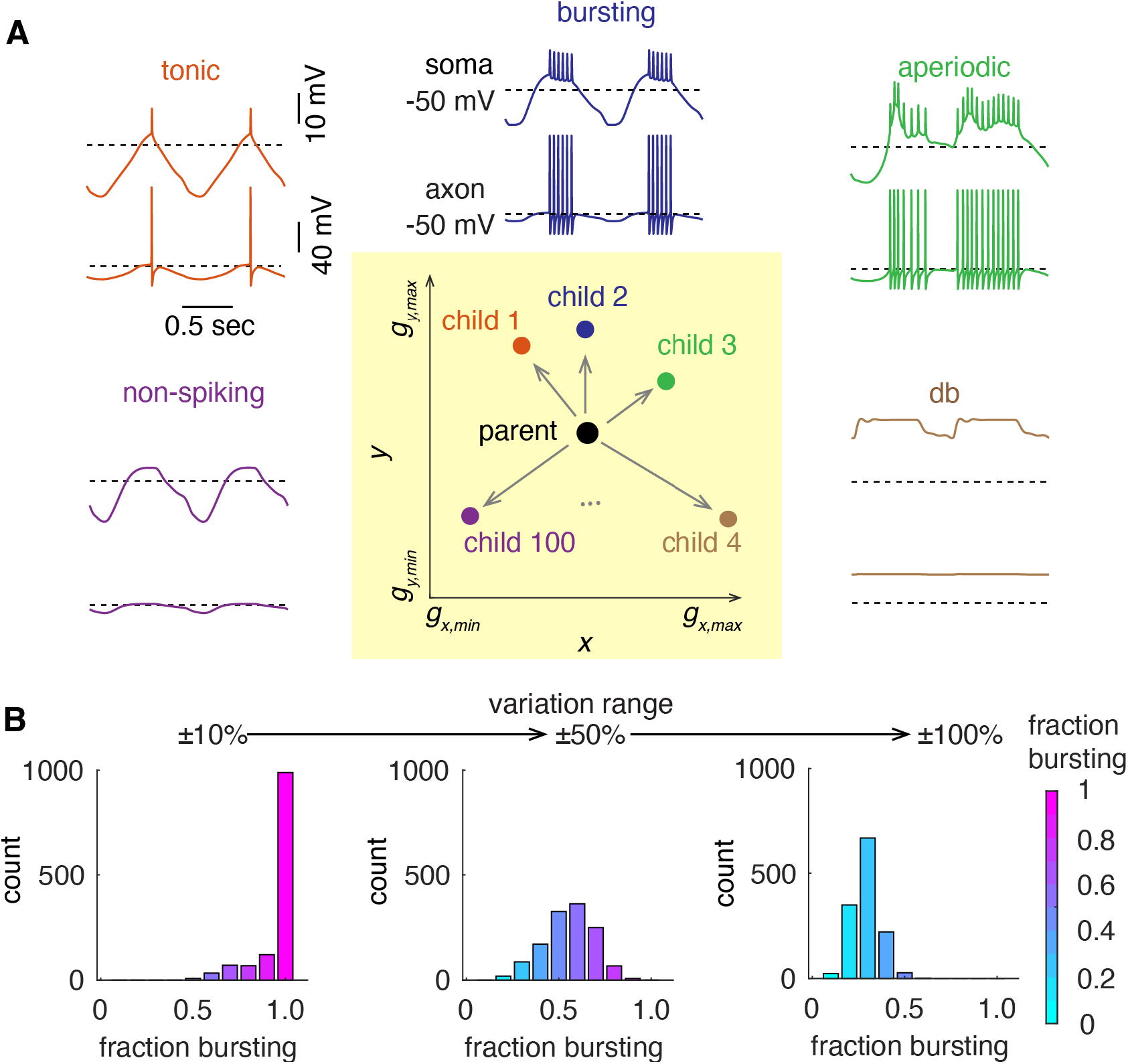
“Noise” perturbation protocol and neuronal resilience to conductance variations. (*A*) 2-D schematic of the channel conductance perturbation protocol (bottom middle, shaded in yellow). *x* and *y* represent different types of ion channels. In each trial, all channel conductances of a “parent” model were simultaneously varied by random values (details in Methods). Each “parent” model (n = 1292) generated new m “child” models after m-trial perturbations. Example firing patterns in “child” models include tonic spiking (child 1), bursting (child 2), aperiodic spiking (child 3), depolarization block (db, child 4), and non-spiking (child 100). (*B*) Histogram of the ratio of “child” models that are bursters for each “parent” model, when models undergo different ranges of random channel conductance variations, ±10%, ±50%, and ±100% from left to right.

The “child” models showed a variety of behaviors, some of which can be generally classified as tonic spiking, rebound bursting, aperiodic spiking, depolarization block or non-spiking (Fig. 2A). The simulation results of 100 trials for each “parent” model are summarized in Figure 2B. For each “parent” model, the ratio of “child” models that are also bursters with at least 2 spikes/burst was calculated to capture its robustness to channel conductance variations. A higher ratio suggests the “parent” model is more robust to channel conductance variations. When channel conductances were simultaneously varied within ±10% of their corresponding parameter ranges, the rebound bursting pattern was remarkably robust to the conductance variations. For 989 of the “parent” models over 90% of the “child” models generated rebound bursting and for 649 of the “parent” models all the “child” models reliably generated rebound bursting. The overall high ratios of “child” models being bursters indicates that the pattern of rebound bursting is relatively insensitive to the conductance variations in these “parent” models. When channel conductances were varied in larger ranges, more and more “parent” models showed decreased ratios of “child” models being bursters, as reflected by their left-shifted distribution in the histograms (Fig. 2B).

It is always interesting to estimate the size and structure of the 14-dimension manifold in which these models exist. Previous work on similar conductance-based models argued that models with similar behavior are in connected regions of parameter space (3, 7). For many “parent” models in these previous simulations (n = 649), the perturbation-generated “child” models reliably generate rebound bursting, suggesting that there are reasonably sized areas around those “parent” models with relatively consistent behaviors. We estimate that the size of the manifold in each channel conductance direction is at least 20% of their corresponding conductance ranges.

Therefore, we selected 37 elite “parent” models whose “child” models reliably generate rebound bursting for 100 trials of perturbations and have 3 – 11 spikes per burst when the variation range is ±10%. Then, we challenged each of them with 10,000 trials of perturbations. For 23 models, all their “child” models reliably generated rebound bursting and in the other 14 models the lowest ratio of the “child” models generating rebound bursting was 98.95%. These results again support the interpretation that this bursting behavior is relatively insensitive to random channel conductance variations. We plotted the spike waveforms and channel conductances for 8 of the 23 models in Figure 3. Except for larger values of the PY neuron synaptic input, the other channel conductance values spread broadly within their corresponding conductance ranges, suggesting that robust models exist widely in the 14-dimension parameter space.

**Fig. 3.**
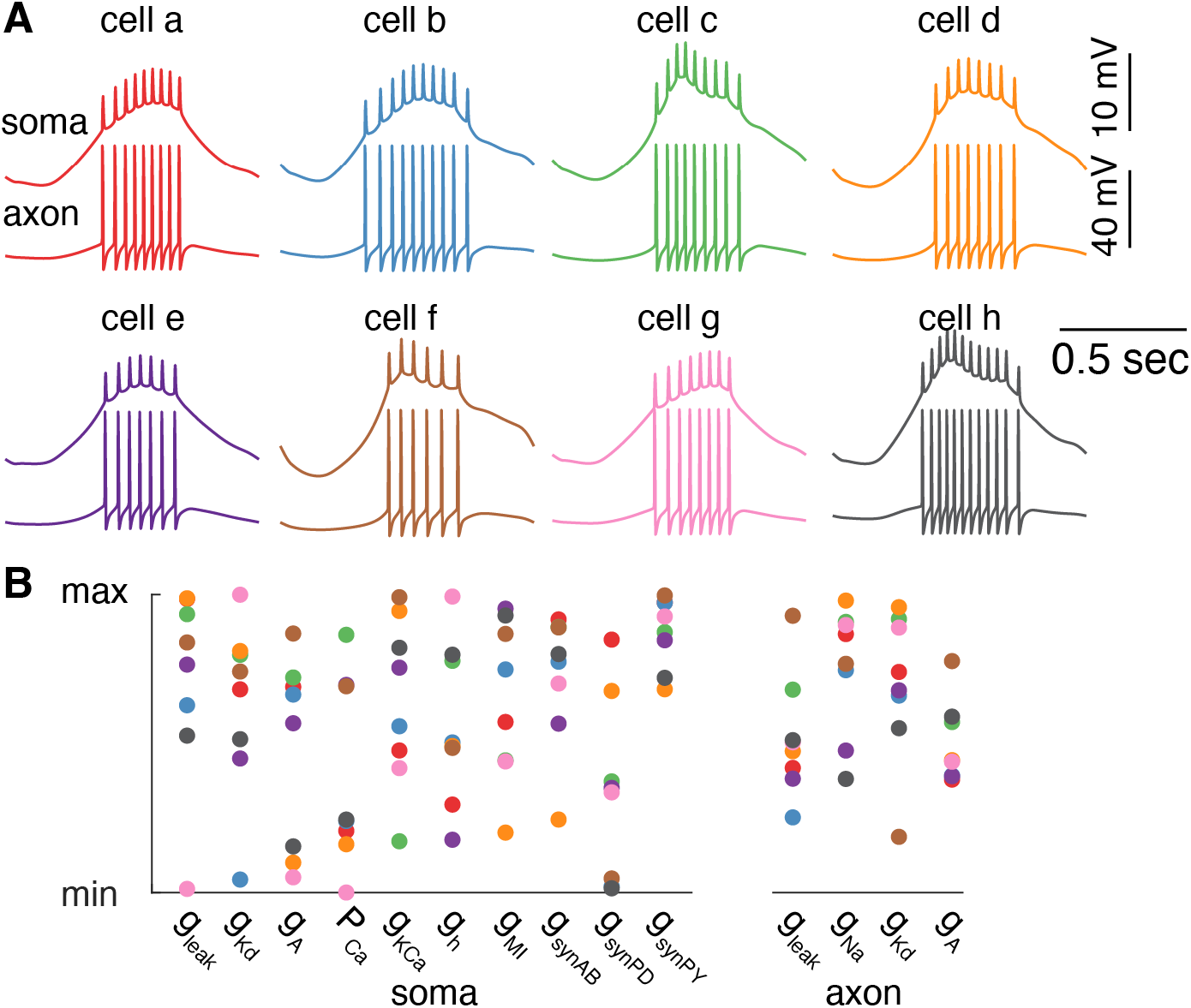
Robust models are widely distributed in parameter space. (*A*) The spike waveforms in the soma (top) and axon (bottom) of 8 example “parent” models (models a – h), whose “child” models reliably produce rebound bursting after 10,000-trial perturbations when the variation range is ±10%. (*B*) The parameter distribution of these 8 example models. Colors code different models (models a – h from panel A).

If one group of neurons shows an advantage in maintaining rebound bursting relative to another group within a small variation range, is this advantage preserved with larger variations? When the variation range increases, the two groups of neurons have similar ratios of maintaining the bursting pattern regardless of their ratios with smaller variation ranges (Fig. 4A).

**Fig. 4.**
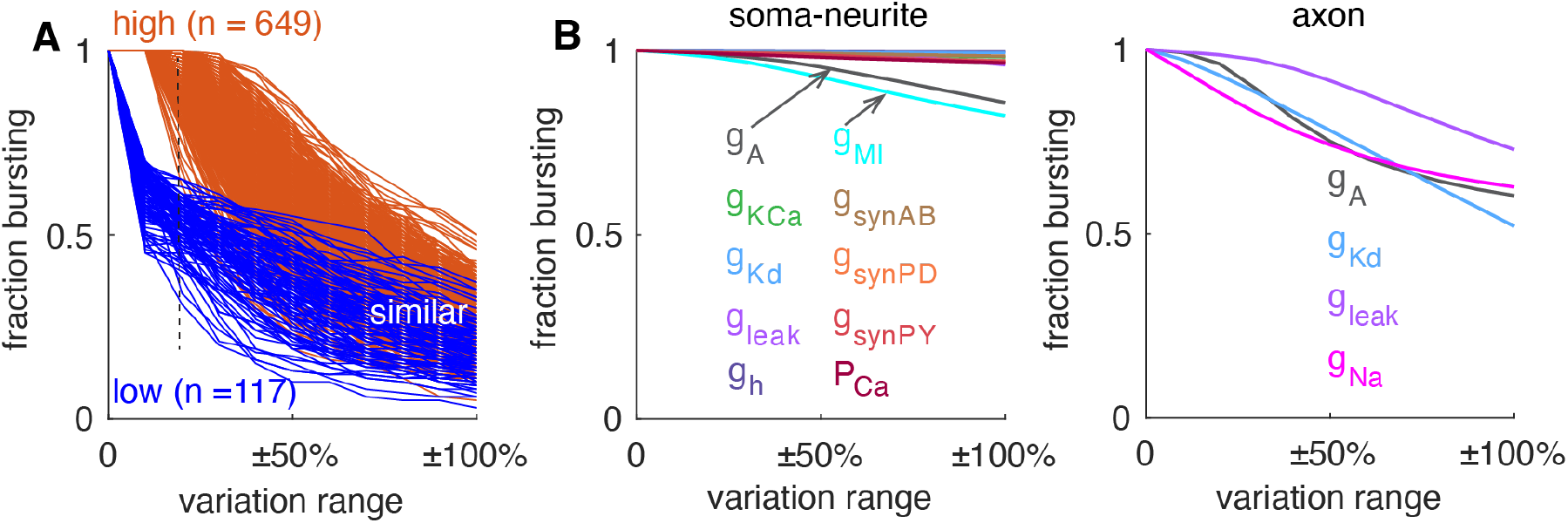
Factors regulating neuronal robustness. (*A*) The effect of variation range on neuronal resilience between two groups of neurons. They are grouped by the ratio of “child” models that are bursters when the variation range is ±10% (orange group, 100%; blue group, <70%). (*B*) The sensitivity of neuronal resilience to individual channel conductance variations (coded by colors). Left and right plots show the effect of varying one of the channel conductances in the soma-neurite and the axon, respectively. The x-axis represents conductance variation range, and the y-axis represents the average ratio of “child” models that are bursters.

We also explored the effect of varying individual channel conductance on the robustness of rebound bursting. Compared with the effect of varying 14 channel conductances simultaneously, the models are more resilient to single channel conductance variations. In the soma and neurites, the model’s performance shows modest sensitivity to I_A_ and I_MI_ conductances. In contrast, varying each of the conductances in the axon changes the model’s firing pattern more efficiently.

### The Effect of Soma-axon Coupling on Axonal Spiking during Bursts

In the LP neuron, the soma and neurites generate rebound slow waves, that evoke bursts of Na^+^ spikes in the axon. Therefore, the soma-axon coupling is key to regulating these bursts of spikes. In the whole model database (n = 1292), we varied the axial resistivity, Ra, between the soma and the axon to change their degree of coupling, while the Ra values in other compartments were fixed at the control value of 100 Ωcm. When the soma to axon Ra was changed, some models no longer produced rebound bursts (Fig. 5A). All models generated rebound bursting when the Ra was in the range of 95 Ωcm – 125 Ωcm (Fig. 5B). For Ra in the range of 40 Ωcm – 300 Ωcm, 1175 (91%) models generated rebound bursting. These results suggest that the bursting pattern is resilient to Ra changes around the control value of 100 Ωcm. When Ra deviated considerably from the control value, the number of rebound bursting neurons decreased significantly. The models that generate rebound bursting with larger Ra deviations are usually bursters when the Ra deviation is lower. For example, in the range of 2 Ωcm – 100 Ωcm, if a model is a burster with a smaller Ra, it is always a burster when the Ra has a larger value, with rare exceptions for the Ra from 15 Ωcm to 20 Ω cm (2 exceptions) and from 35 Ωcm to 40 Ωcm (1 exception).

**Fig. 5.**
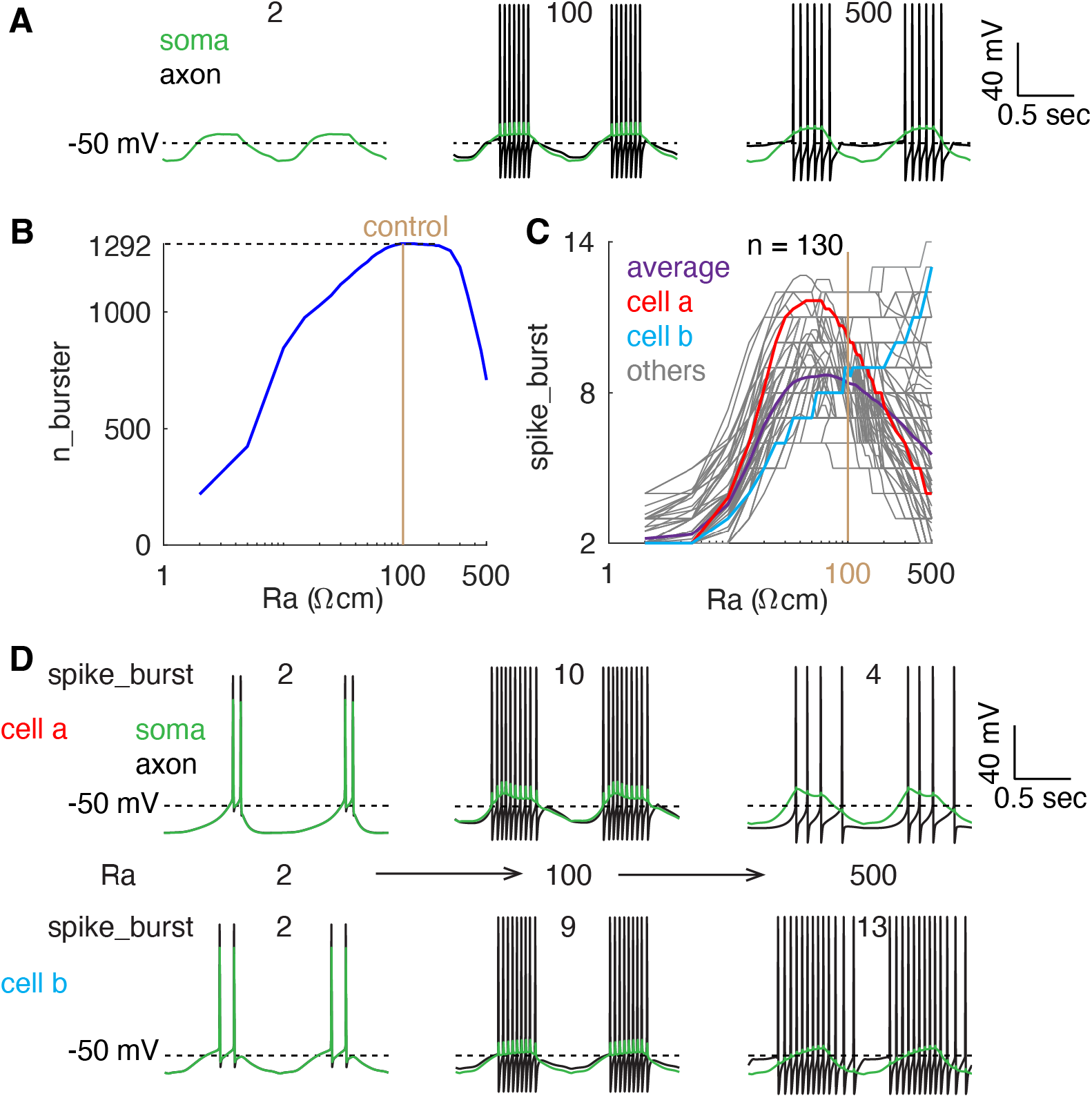
The effect of soma-axon coupling. (*A*) Example somatic (green) and axonal (black) spike waveforms when Ra was changed from 100 Ωcm (middle) to 2 Ωcm (left) and 500 Ωcm (right). (*B*) The relationship between the soma to axon Ra and the bursting neuron population size (n_burster). n = 1292 when Ra = 100 Ωcm under control condition. The x-axis is a logarithmic scale. (*C*) The relationship between the soma to axon Ra and the number of spikes per burst (spike_burst) in neurons that generate rebound bursting regardless of Ra changes (n = 130). Purple, population average; red and cyan, two example neurons; gray, other neurons. The x-axis is a logarithmic scale. (*D*) The effect of soma to axon Ra on somatic (green) and axonal (black) bursting changes in the example cells (cells a and b) highlighted in C (Ra, 2 Ωcm → 500 Ωcm from left to right). In all simulations, there are no channel conductance variations.

The simulation results above show that the degree of soma-axon coupling affects the population size of bursting neurons. We selected 130 models that always generate the pattern of rebound bursting for Ra within 2 Ωcm – 500 Ωcm. The number of spikes per burst reflects the ability of the axon to spike during bursts. For most of these neurons, the number of spikes per burst shows biphasic changes with Ra, as also reflected by the average changes (Fig. 5C).

However, in several neuron models, this number monotonically increases within the tested Ra range.

When Ra is small, the soma and the axon are almost isopotential, as revealed by the nearly overlapped somatic and axonal membrane potentials (Fig. 5D, left). When the soma and axon are strongly coupled, this reduces the ability of the axon to spike during bursts for two reasons. First, the axonal membrane potential after each spike is closer to the somatic slow-wave peak (bottom Vm during the bursts), which reduces the recovery of Na^+^ channels from inactivation. Second, the soma exerts a larger capacitance load on the axon during spiking (top Vm during the bursts).

Therefore, when the Ra was 2 Ωcm, the number of spikes per burst was low. Increasing Ra to make the soma and the axon more isolated should increase the axonal ability to spike during bursts, considering just the above 2 factors. However, the increased isolation between the soma and the axon also reduces the axial current from the somatic slow wave that triggers axonal spiking during bursts. Consequently, changes in the number of spikes per burst depend on the balance of these 3 factors when Ra changes (Fig. 5D).

### The Effect of Soma-axon Coupling on Neuronal Robustness

We explored how soma-axon coupling affects neuronal robustness for all of the 1292 “parent” models. When the soma to axon Ra is near the control value of 100 Ωcm, the average ratio of “child” models that are bursters is insensitive to Ra changes irrespective of conductance variation ranges (Fig. 6A1, Ra = 50, 100 and 125 Ωcm). However, when the soma-axon coupling gets too strong (Ra = 2 Ωcm) or too weak (Ra = 500 Ωcm), the average ratio of “child” models that are bursters is significantly reduced. Under these conditions, for many “parent” models that lose rebound bursting, their “child” models become bursters when the channel conductances are perturbed to appropriate ranges (Fig. 6A2, A3). When Ra is 2 Ωcm, the average ratio of “child” models that are bursters shows biphasic changes. Under these conditions, some of the original “parent” models are not bursting neurons, but in the neighboring parameter space they may remain bursting (Fig. 6A2, A3). Figure 6 suggests that the “density” of bursting neuron models in the whole parameter space is nearly unchanged when Ra is near 100 Ωcm but significantly decreases when Ra deviates further from its control value.

**Fig. 6.**
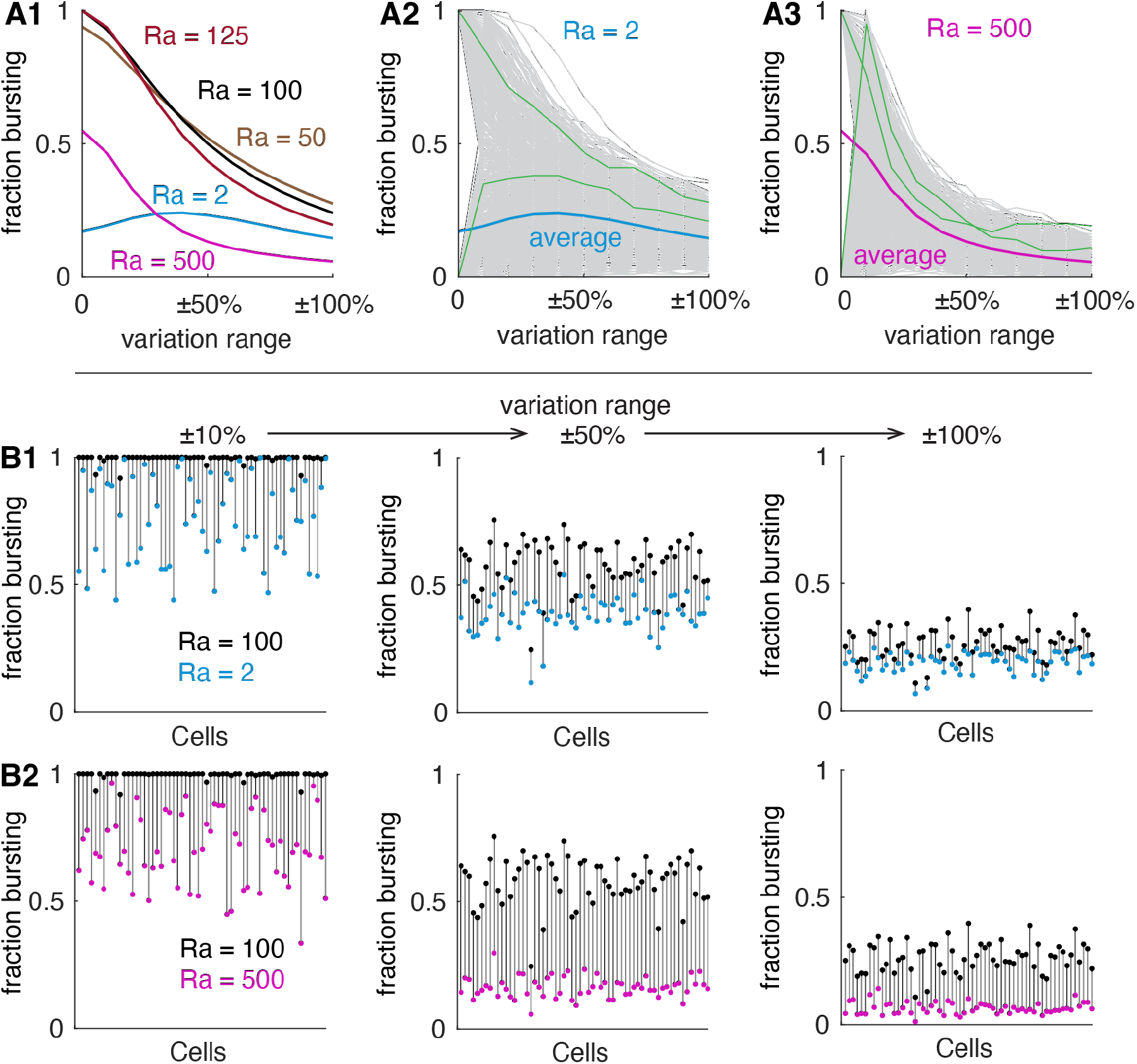
The effect of soma-axon coupling on neuronal resilience to conductance variations. (*A*) The effect of soma to axon Ra on neuronal resilience to conductance variations for all the “parent” models. A1, The average ratio of “child” models that are bursters with different Ra values (coded by colors). A2 and A3 show individual neuronal ratios of “child” models that are bursters for Ra = 2 Ωcm and 500 Ωcm, respectively. Green traces show typical example plots. Blue and magenta traces show population average ratios, as in A1. Gray traces show other individual neurons. (*B*) The comparison of each neuronal ratio of “child” models that are bursters when Ra is changed from the control value of 100 Ωcm. B1, Ra is changed from 100 Ωcm (black dots) to 2 Ωcm (blue dots); B2, Ra is changed from 100 Ωcm (black dots) to 500 Ωcm (magenta dots). From left to right, conductance variation range increases from ±10% to ±100%. Eighty of the total 131 elite “parent” models are shown.

To explore how the degree of soma-axon coupling affects the ability of individual neurons to maintain rebound bursting when faced with channel conductance variations, we selected 131 elite models from the 1292 “parent” models. These 131 elite “parent” models reliably generate rebound bursting when the soma to axon Ra is 2 Ωcm, 100 Ωcm, and 500 Ωcm. Strikingly, for each of these “parent” models, the ratio of “child” models that are bursters consistently decreases when Ra is changed from 100 Ωcm to 2 Ωcm and from 100 Ωcm to 500 Ωcm (Fig. 6B, only 80 cells were plotted, but all cells show the same trend), regardless of the conductance variation ranges. These simulation results suggest that the degree of soma-axon coupling determines the size of the manifold that corresponds to bursting neurons and therefore the neuronal robustness to channel conductance variations.

## Discussion

Most neurons have complex structures over which ion channel distributions must be established such that the neuron’s characteristic electrical properties are maintained, given the inevitable “noise” that must accompany the ongoing turnover of channels in the membrane. It is known that some neurons display specific patterns of channel distributions that are thought to aid in the neuronal computations that these neurons perform (25-27). Therefore, it becomes important to determine the extent to which the spatial structure of neurons makes them more or less sensitive to fluctuations in channel densities across their structures.

In this work, we used a population of LP neuron compartmental models developed previously (5) to explore neuronal robustness to channel conductance variations. All members of this population met a series of criteria that ensure that they capture many of the essential features of the biological neurons whose behavior they were intended to capture (5). At the same time, these models show considerable degeneracy in that the specific values of their conductances are highly variable (5). While these models separate the soma, axon, and dendritic compartments, and are therefore more biologically “realistic” than single compartment, point, neurons (28), they are not intended to capture all of the detailed structures of individual STG neurons.

In models of one or two dimensions, it is easy to find the parameter sets that define the boundary between different types of neuronal activity patterns (29, 30). However, as the number of dimensions increases, it becomes more challenging to characterize the parameter space for neurons with specific properties (7, 16). For the 14-dimension parameter space here, it would be a daunting task to locate all possible bursting models with a grid search, because it is impossible to know whether the sampling step size is small enough (3). Instead, we used random parameter perturbations. This approach does not provide an unambiguous map of the entire manifold.

Nonetheless, if all “child” models are bursters after exerting random and simultaneous conductance perturbations for sufficiently many trials, it argues the existence of a multi-dimension manifold that contains the very large majority of models of similar behavior (7). We did this in a subset of neuron models for 10,000 trials and most of them reliably generated rebound bursting after perturbations (Fig. 3).

Multiple mechanisms likely exist that can allow neurons to compensate for perturbation-caused changes. In homeostatic models of intrinsic excitability and regulation of synaptic strength, both ion channel conductances and synaptic strength can be altered to recover neuronal and circuit function properties in response to deviations from a set-point (14, 31-36).

Some homeostatic mechanisms require new protein synthesis, and therefore recovery may be slow and require hours or even days, whereas other processes may be more rapid (33, 35, 37, 38).

Neurons constantly face diverse perturbations such as temperature changes, neuromodulation, and stochastic changes in protein synthesis, but they rarely fail in healthy animals under physiological conditions. This argues that most neurons are not sitting too close to bifurcations, but neurons and circuits can tolerate some changes in their parameters without loss of function. The effect of morphology on the stability of physiological activity has received less attention than it probably merits. Nonetheless, there are examples of structural alterations that can be homeostatically regulated to control activity (39).

In many neurons, including those of the STG, neuronal Na^+^ spikes are initiated in the axon. In STG neurons, the slow waves that underlie bursting depend on channels in the soma and major neurites; Synaptic inputs and outputs occur on the processes, and these various regions of the neuron are coupled appropriately (5). Biological ranges of soma-axon coupling can increase the density and the size of manifolds that allow rebound bursting (Figs. 5, 6). Recent data argue that the detailed morphology of STG neurons are not important for their overall function (21-23). Thus, instead of requiring the detailed specification of the number and location of each ion channel, this strategy requires only the specification of a generic structure, that in turn depends on an appropriate range of values of the electrical coupling between the soma and the axon. The control value of 100 Ωcm for the Ra in this model falls in the middle range of experimentally reported values (40) and was originally chosen to make the voltage waveforms of the LP neuron model roughly match experimental data (5). While individual LP and other STG neurons show substantial animal-to-animal variability in channel conductances and channel mRNAs (1, 41-46), it is important to remember that the motor patterns produced by these neurons in the pyloric circuit are tightly maintained, and the LP neuron’s rebound burst is critical for its proper function (24).

The implications in this work amplify those of the sloppy morphology observed in crab stomatogastric ganglion neurons (21-23). Although the morphological structures of these neurons exhibit significant cell-to-cell variations, they are electrotonically compact with gradually tapering neurites from the soma to the distal tips. This elegant, but not too rigid, solution makes graded transmission resilient to electrotonic decrement across sparsely distributed, synchronous presynaptic sites. The separation of synaptic integration and slow waves from axonal spike initiation zones may be a morphological strategy established to avoid shunting of synaptic currents (21-23, 47). Here, we use models to demonstrate that this division of labor can make neurons more robust to perturbations.

The findings in this study may be relevant to other neuronal systems. Spike bursts are widely observed in the cerebellum (48, 49), the neocortex (50), and the hippocampus (51). For example, the inferior olive neurons in the cerebellum show significant similarities to the LP neuron. Inferior olive neurons show subthreshold oscillations in the soma and then trigger axonal spike bursts to guide cerebellum-related movement and learning (48). Additionally, soma-axon coupling is also critical for the initiation of axonal spike bursts (complex spikes) in cerebellar Purkinje cells (15, 52). Because of the functional importance of these signals in cerebellar coding and learning, the reliability of triggering axonal bursting determines the robustness of corresponding behavioral functions. Moreover, neurons may utilize their entire structure to maintain a robust functional output in other spiking patterns (53, 54).

## Materials and Methods

We used the LP neuron multicompartment models constructed previously (5) and implemented them in NEURON 7.6 (55). These models were constructed to replicate the rebound bursting properties observed in the LP neuron of the crab stomatogastric ganglion. We adopted 1292 out of the original 1304 cell models. The excluded cell models either need a long time to generate rebound bursting or didn’t generate this pattern after reaching a steady state, possibly because of NEURON version changes.

The current types and compartmental distributions are shown in Fig. 1. The detailed mathematical description of these currents can be found in the original work (5). All current models were models as ohmic currents, except I_Ca_. For the ohmic current, 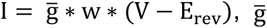 is the maximal conductance, w is the product of activation and inactivation gate state variables, V is the transmembrane potential and E_rev_ is the reversal potential. The calcium current was modeled by Goldman-Hodgkin-Katz equations, 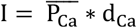, where 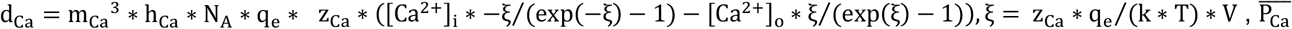 is the maximal permeability of the calcium ions, m_Ca_is the calcium channel activation gate, h_Ca_is the calcium channel inactivation gate, N_A_ is the Avogadro’s constant, q_e_ is the elementary charge, z_Ca_ is the valence of a calcium ion, [Ca^2+^]_i_ and [Ca^2+^]_o_ are intracellular and extracellular calcium concentrations, k is the Boltzmann’s constant and T is the Kelvin temperature. The three types of inhibitory synaptic inputs were approximated by fixed rhythmic patterns of synaptic conductances. For each type of synapse, the synaptic conductance was approximated by 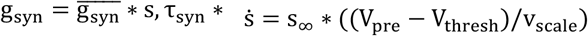, where 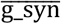 is the maximal synaptic conductance, s is the fraction of synaptic activation, τ_syn_ is the time constant, V_pre_ is the presynaptic voltage (smoothed experimental recordings), V_thresh_ is the synaptic threshold, and v_scale_ is the voltage sensitivity of the synaptic activation. The AB neuron and PY neuron synapses have instantaneous dynamics and a reversal potential of -70 mV. The PD neuron synapse has slower dynamics (with a fixed time constant of 50 ms) and a reversal potential of -80 mV, in keeping with the known properties of these synapses (56).

For conductance perturbation simulations within a variation range, all 14 channel conductances were simultaneously and independently varied by values randomly sampled in the range of [-1, +1]*variation_range*(g_*x,max*_ - g_*x,min*_). *x* represents a type of ionic current; g_*x,max*_ and g_*x,min*_ represent the maximal and the minimal conductances of current *x* in the parameter space, respectively. A conductance was set to zero if the value became negative after a random perturbation. A “child” model was classified as a burster if it fires periodically and there were at least 2 spikes per burst. For the elite “parent” models in Fig. 3, the first criterion is all their “child” models need to be bursters when the variation range is ±10%. The second criterion is the number of spikes per burst in the “parent” models and their corresponding “child” models need to be in the range of 3 – 11 (5). For the results in Figs. 5 and 6, the axial resistivity Ra between the soma and the axon was varied to simulate different degrees of soma-axon coupling. We constrained the Ra within the range of 2 Ωcm – 500 Ωcm, to maintain a sufficient number of neurons in Fig. 5C and Fig. 6B to compare the changes in individual neurons.

## Data Availability

Model code will be available in ModelDB.

## Acknowledgments

This work was supported by NIH Grants R35NS097343 and R01MH046742. Y.Z. would like to thank Adam Taylor for sharing the model code and Sonal Kedia for sharing experimental data.

